# NOTCH1 drives immune-escape mechanisms in B cell malignancies

**DOI:** 10.1101/2021.04.10.439192

**Authors:** Maurizio Mangolini, Alba Maiques-Diaz, Stella Charalampopoulou, Elena Gerhard-Hartmann, Johannes Bloehdorn, Andrew Moore, Junyan Lu, Valar Nila Roamio Franklin, Chandra Sekkar Reddy Chilamakuri, Ilias Moutsoupoulos, Andreas Rosenwald, Stephan Stilgenbauer, Thorsten Zenz, Irina Mohorianu, Clive D’Santos, Silvia Deaglio, Jose I. Martin-Subero, Ingo Ringshausen

## Abstract

*NOTCH1* is a recurrently mutated gene in Chronic Lymphocytic Leukemia (CLL) and Mantle Cell Lymphoma (MCL). Functional studies to investigate its role have been hampered by the inability to genetically manipulate primary human lymphoma cells, attributed to low transduction-efficacy and procedure-associated toxicity. To overcome these limitations, we have developed a novel method to retrovirally transfer genes into malignant human B cells. We generated isogenic human tumor cells from patients with CLL and MCL, differing only in their expression of NOTCH1. Our data demonstrate that NOTCH1 facilitates immune-escape of malignant B cells by up-regulating PD-L1, partly dependent on autocrine interferon-γ signaling. In addition, NOTCH1 causes silencing of the entire HLA-class II locus via suppression of the transcriptional co-activator CIITA. These NOTCH1-mediated immune escape mechanisms are associated with the expansion of CD4^+^ T cells *in vivo*, further contributing to the poor clinical outcome of *NOTCH1*-mutated CLL and MCL.

## Introduction

The landscape of somatic mutations present in malignant B cells from patients with CLL has been described in several pivotal sequencing studies, identifying more than 40 recurrent mutations mostly affecting oncogenes (*NOTCH1*, Wnt-signalling), tumor suppressors (*TP53, ATM*) and genes involved in RNA-processing (*SF3B1, XPO1, RPS15*)^1–4^. While the prognostic significance is known for some of these mutations, their specific contributions to the pathogenesis of the disease remains largely unknown. Attempts to address this experimentally have employed genetically engineered mouse models (GEMMs), most commonly the Eμ-TCL1 mouse model^5^, and human cell lines. While the former model proved to be useful in particular to recapitulate disease aspects which can only properly be studied *in vivo* (such as tumor-microenvironment interactions), significant limitations exist which prevent extrapolation of data from mice to human. In addition, the experimental manipulation of primary tumor cells from the Eμ-TCL1-model remains technically challenging and most commonly requires crossing of different GEMMs, which consumes time and resources. In contrast, cell lines are easy to manipulate and provide a sheer unlimited and immediate access to tumor cells. However, because they are commonly obtained from patients with end-stage, refractory diseases, cell lines frequently are EBV-positive^6,7^ and often have been selected for decades to grow in minimal culture conditions, therefore losing the biological identity they are supposed to represent. Vigorous proliferation, absence of spontaneous apoptosis and aberrant homing in NSG mice are some examples for these discrepancies, limiting the conclusions one can draw from such experiments.

Few studies have used adenovirus vectors or their derivates to genetically manipulate primary CLL cells^8^. However, the lack of integration into the host genome results in only transient expression of a gene-of-interest (GOI) and precludes from studying effects in dividing cells or subsequent use of cells in *in vivo* studies. Alternative attempts using retro- or lentivirus vectors for gene transfer have been unsuccessful for decades, limited by low transfection efficacy (<1%) and substantial toxicity^9^, which made down-stream analyses impossible. To overcome these limitations, we have developed a novel method to effectively infect primary neoplastic human B cells. This method permits gene transfer with high transduction efficacy and minimal toxicity, which allows the functional investigations of genes recurrently mutated in primary malignant B cells. We used patient samples from CLL and MCL, two mature B neoplasms with partially overlapping biological features and clinical behaviors^10^ that lack appropriate *in vitro* or *in vivo* models that span their clinico-biological spectrum.

We have employed this technique to interrogate the molecular functions of NOTCH1, which is one of the most commonly mutated genes in CLL and associated with a poor clinical outcome and a high frequency of Richter’s transformation^11–13^. Most NOTCH1-activating mutations affect the PEST-domain, encoded by exon 34^14^. In addition, point mutations in the 3’-UTR have been identified and cause expression of a truncated protein^2^. Both scenarios result in an abnormally stable NOTCH1 protein, which continues to require ligand-binding in order to become transcriptionally active. Several groups have employed cell lines to study the biology of NOTCH1 in B cell malignancies and then associated these findings with data from primary cells^15–18^. While such approaches provided important insights into the role of NOTCH1 in CLL, it often remains unclear whether these findings report a direct consequence of activated NOTCH1 or are a mere correlation. Our method to retrovirally infect primary malignant B cells from patients with CLL and MCL to generate isogenic cells provides a unique opportunity to answer this question. Here we provide evidence of how NOTCH1 favors immune escape of tumor B cells and we address how cells with trisomy 12 may provide a selective advantage for *NOTCH1*-mutations.

## Results

### Retroviral transduction into primary human tumor B cells permits functional downstream analyses

In the past, transduction of CLL B cells using retroviral or lentiviral vectors has largely been unsuccessful mainly due to the fact that cells *ex vivo* are prone to apoptosis and are arrested in the G0/G1 phase of the cell cycle^9^. In order to overcome these limitations, we generated three different *de-novo* retrovirus-based envelopes able to infect mammalian cells of different tissue origins. Additionally, we engineered a new stroma-cell line from a subset of CD45^neg^, Lin^neg^, Sca1^pos^ -bone marrow derived stromal cells, hereafter called MM1-cells. We initially tested different viral envelopes to genetically manipulate proliferating CLL cells, which were continuously cultured on MM1-cells after transduction. Initial attempts to transduce cells were unsuccessful due to a persistent high rate of apoptosis of also proliferative cells and required further manipulation of MM1-cells. MM1-cells constitutively expressing 3 additional prosurvival factors fully antagonized procedure-associated toxicity and permitted the successful transduction of primary CLL cells. While primary malignant B cells were effectively transduced with Env1 (24% in CLL and 12% in MCL), the Env2 infected a higher percentage of CLL and MCL cells with an average efficacy of 37% and 39%, respectively. In contrast, CLL cells were resistant to infections with the Env3, which displayed moderate efficacy to infect primary MCL cells (Fig. 1a). Importantly, our transfection method was not associated with increased cell death: after 7 days, CLL cell viability was >80% (Fig. 1b), similar to non-transduced, previously cryopreserved cells and cultured under identical conditions (Supplementary Fig. 1a).

**Figure 1:**
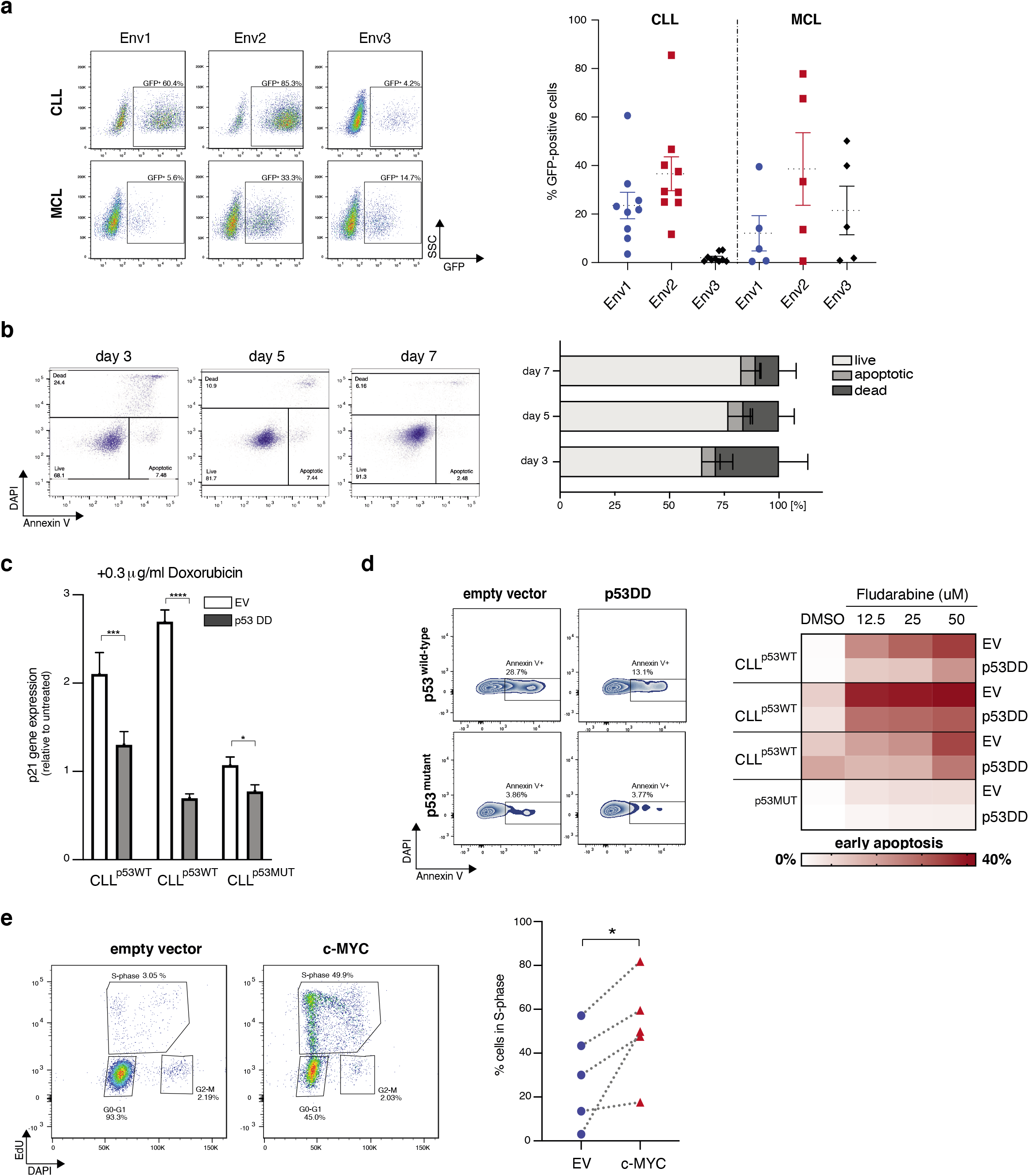
Viral transduction of primary human B cells from CLL/ MCL patients. a. Quantification of transduction efficiency as determined by GFP expression using three different envelope constructs in CLL and MCL primary cells 72h post-transduction. Representative flow cytometry images of CLL and MCL transduction efficacy are shown for each construct on the left. b. Quantification of viable (DAPI^-^ Annexin V^-^), apoptotic (DAPI^-^ Annexin-V^+^) and dead (DAPI^+^) cells 3-, 5- and 7-days post transduction (*n=8*). Representative flow cytometry images and gating strategy are shown for each time point. c. qRT-PCR analysis of *p21* mRNA expression in primary CLL cells transduced with an empty vector control (EV) or a dominant negative P53 expressing vector (P53DD) after 12h treatment with doxorubicin, normalized to cells treated with a vehicle control (DMSO). d. Heatmap showing the percentage of apoptotic (DAPI^-^Annexin-V^+^) CLL cells transduced with either an empty control (EV) or a dominant negative P53 expressing vector (P53DD) after 24h exposure to increasing concentrations of Fludarabine. Annexin-V positivity was assessed on GFP only expressing cells. Representative flow cytometry images are shown for each drug concentration on the left. e. S-phase quantification of CLL cells transduced with an empty vector or a MYC expressing vector 6 days post transduction (*n=5*). Cells co-cultured with feeder cells were pulsed with Edu for 12h before fixation. Cohorts are shown as mean ± SD. *P < 0.05, **P < 0.01, ***P < 0.001, ****P < 0.0001 and not significant (ns) P > 0.05.

To test that transduced cells remained functional to allow down-stream analyses, cells were transduced with a dominant-negative TP53 (p53DD) and subsequently exposed to the anthracycline doxorubicin for 12 hours. P53DD significantly mitigated *p21*mRNA transcription induced by doxorubicin in cells carrying wild-type *TP53*, but significantly less so in p53-deficient cells (Fig. 1c). Similarly, p53DD reduced Fludarabine-induced apoptosis (Fig. 1d). In addition to the interference with a major tumor suppressor, we overexpressed the proto-oncogene c-MYC in primary malignant B cells from CLL patients. Expectedly, ectopically expressed c-MYC induced a robust proliferative response in CLL cells, indicated by the increased number of cells transitioning through S-phase (Fig. 1e).

In conclusion, we have established a method to effectively infect primary tumor cells from patients with CLL and MCL with minimal toxicity, allowing to generate isogenic, patient specific tumor cells which differ only in the GOI.

### NOTCH1 drives proliferation, CD38-expression and enhances B cell receptor signaling

We next used this method to investigate the role of NOTCH1 in primary CLL cells, which is frequently mutated in approximately 10% of untreated patients^14,19^. In order to simultaneously account for point-mutations, missense and frameshift mutations affecting exon 34 and for less common 3’-UTR *NOTCH1* mutations, we expressed the coding sequence of NOTCH1-ICD, but lacking the PEST-domain, followed by IRES-GFP (hereafter named *NOTCH1^ΔPEST^*; Fig. 2a) in primary CLL cells. Importantly, since NOTCH1^ΔPEST^ also lacked the ectodomain, activation did not require binding of NOTCH ligands or cleavage of the extracellular domain for transactivation. Since CLL cells co-express NOTCH1 and NOTCH2^20^, this approach allowed us to investigate the function of NOTCH1 in isolation without simultaneously activating other NOTCH-receptors. To test that NOTCH1^ΔPEST^ was transcriptionally active, we first assessed the mRNA expression levels of *HES1, HEY1* and *DTX1*, bona-fide NOTCH-target genes. Compared to empty vector (EV) control, NOTCH1^ΔPEST^ (N1^ΔPEST^) increased the abundancy of *HES1, HEY1* and *DTX1* mRNA by 3.0-, 3.4- and 3.6-fold, respectively (Fig. 2b). Importantly, similar expression changes of *HES1* and *DTX1* were reported in ligand activated CLL cells carrying *NOTCH1* mutations^17,18,21,22^, indicating that NOTCH1^ΔPEST^ has a similar activation potential than mutated, endogenous NOTCH1.

**Figure 2:**
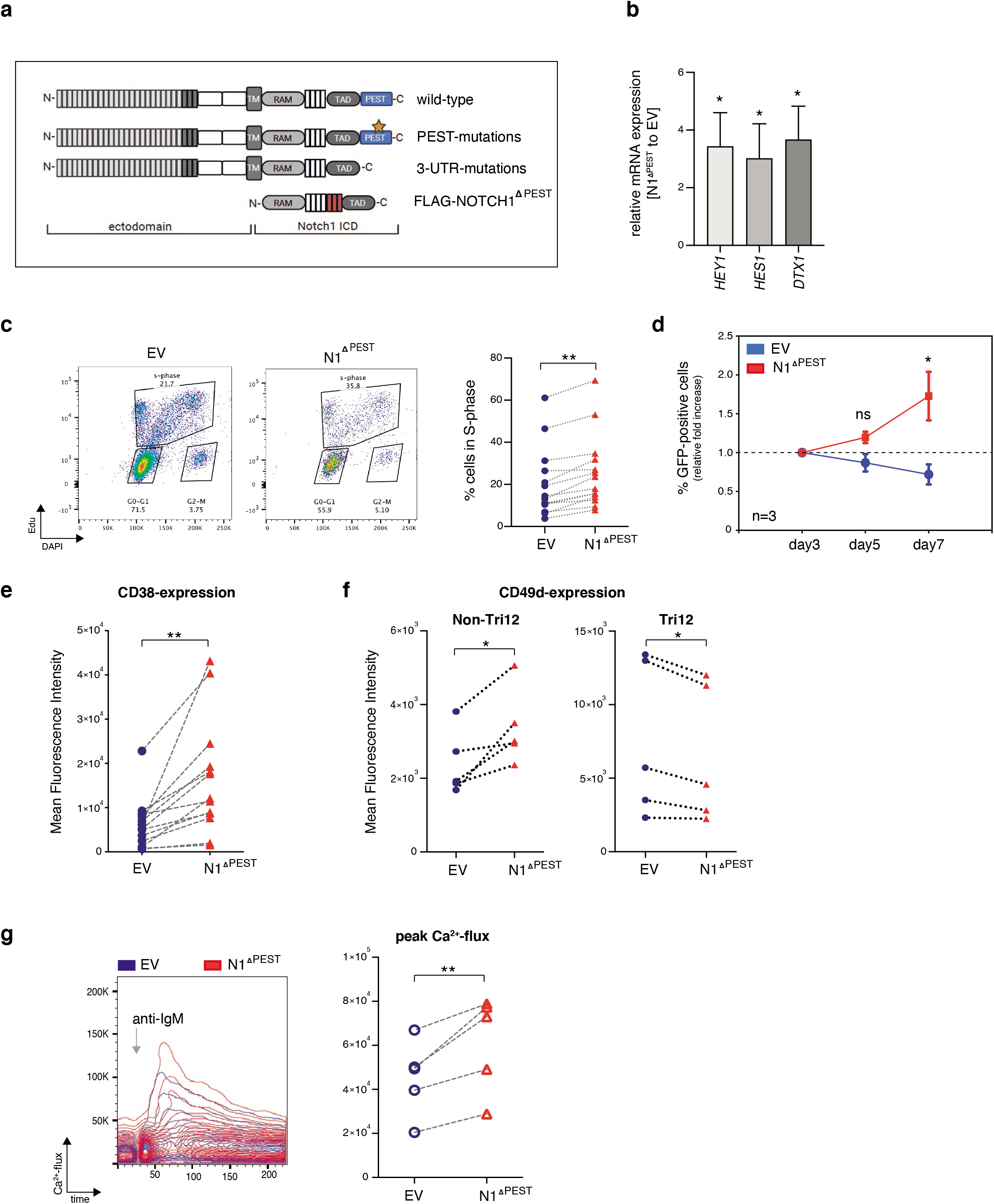
NOTCH1^ΔPEST^ recapitulates biological features of mutated NOTCH1. a. Schematic representation of recurrent NOTCH1 mutations and the NOTCH1^ΔPEST^ construct. b. qRT-PCR analysis of canonical NOTCH1^ΔPEST^ target genes expression (*HEY1, HES1, DTX1*) in primary CLL cells transduced with *NOTCH1^ΔPEST^*. Expression is normalized to cells transduced with an empty vector control (*n=5*). Cells were FACS sorted for GFP expression prior to RNA extraction 5 days post transduction. Error bars are shown as mean ± SD. c. S-phase quantification of CLL cells transduced with an *empty vector* (blue) or *NOTCH1^ΔPEST^* (red) (*n=13*). Cells co-cultured on feeder cells were pulsed with Edu for 12h before fixation. Cell cycle analysis was restricted to GFP-positive cells; representative flow cytometry images are shown on the left. d. Growth curve of CLL cells transduced with an *empty vector* (blue line) or *NOTCH1^ΔPEST^* (red line) quantified as GFP expression 3-, 5- and 7-days post transduction. The mean percentage of GFP-positive cells (+/-SD) of 3 independent patients is shown. e. CD38 expression on CLL cells transduced with an *empty vector* (blue) or *NOTCH1^ΔPEST^* (red) 5 days post transduction (*n=13*), assessed by flow cytometry. Representative flow cytometry images are shown on the right. f. CD49d expression on CLL cells (del13q (*n=5*, left) or trisomy 12 (*n=5*, right) transduced with an *empty vector* (blue) or *NOTCH1^ΔPEST^* (red) 5 days post transduction. g. Quantification of peak of Ca^2+^-flux in interval time in response to anti-IgM stimulation of CLL cells expressing NOTCH1^ΔPEST^ (red) identified by GFP expression compared to GFP negative cells (blue) (*n=5*). Interval times were automatically determined by kinetic analysis using the FlowJo software. Representative Ca^2+^-flux dot plots overlap of a single patient is shown on the left showing the induction of Ca^2+^-flux following IgM stimulation. *P < 0.05, **P < 0.01, ***P < 0.001, ****P < 0.0001 and not significant (ns) P > 0.05.

Several previous studies have suggested that CLL cells carrying *NOTCH1* mutations have a proliferative advantage compared to wild-type CLL cells^16,22^. To test whether NOTCH1 is involved in cell cycle regulation, we assessed the number of GFP^+^ cells in S-phase 6 days after transduction. Compared to *EV*-control cells, primary CLL cells expressing *NOTCH1^ΔPEST^* consistently showed a higher percentage of cells going through S-phase (Fig. 2c). Accordingly, the fraction of GFP-positive CLL cells continuously increased only in the *NOTCH1^ΔPEST^*-transduced cells but not in *EV*-controls, indicating that NOTCH1 positively affects cell cycle progression (Fig. 2d).

*NOTCH1* mutations have also been associated with surface CD38 expression^23^. In keeping with the observation that CD38 expression is higher in CLL lymph nodes compared to peripheral blood cells^24^, co-culture of CLL cells on MM1-cells increased baseline expression of CD38 (data not shown). NOTCH1^ΔPEST^ consistently up-regulated CD38 expression further, suggesting that its expression is functionally dependent on NOTCH1 activation (Fig. 2e). Of note, NOTCH1^ΔPEST^ expression did not induce the expression of CD138, *BLIMP1* or *IRF4* (Supplementary Fig. 1b,c), indicating that CLL B cells did not differentiate into antibody-secreting plasma cells ^25^. Other studies have associated NOTCH1 mutations to the expression of CD49d, which equally and independently indicates a poor prognosis. In agreement with a previous study on MEC-1 cells^18^, we observed that NOTCH1^ΔPEST^ induced surface expression of CD49d only in non-trisomy 12 patients, which overall expressed much higher levels of CD49d (Fig. 2f). Lastly, we assessed B cell receptor (BCR)-responsiveness of CLL cells transduced with *NOTCH1^ΔPEST^* or *EV*. Anti-IgM induced a stronger calcium-flux in cells expressing *NOTCH1^ΔPEST^* compared to *EV* cells (Fig. 2g), supporting a recent study which demonstrated collaboration between NOTCH- and BCR-signaling ^21^. In conclusion, retrovirally expressed *NOTCH1^ΔPEST^* has biological activities similar to mutated NOTCH1 and recapitulates functions previously attributed to NOTCH-activation.

### Patients with trisomy-12 or del13q present a common NOTCH1 transcriptome

To define the global transcriptional program controlled by NOTCH1 and contributing to these phenotypic and proliferative effects, we performed RNA-seq on 13 primary CLL samples, either transduced with an *EV*-control *or NOTCH1^ΔPEST^*. Only patients who had a deletion of chromosome 13q (del13q) or carried an additional copy of chromosome 12 (tri12), assessed by conventional FISH-analyses, were included. Pairwise analysis of all 13 patient samples identified 1636 differentially expressed genes of which 979 were upregulated and 657 were downregulated (applying a cut-off of Log2FC>0.5, present in at least half of all samples) (Fig. 3a). Gene Set Enrichment Analysis (GSEA) of these differentially expressed (DE) genes identified gene clusters in canonical Notch-signaling (e.g., *HES4, SEMA7A, CD300A* and *DTX1*), B cell activation/ BCR-signaling (e.g., *FYN, BLNK* and *CR2*) and MAPK-activation (e.g., *MAPK8* and *MAP2K6*), in keeping with previous reports based on the ectopic expression of NOTCH1 in lymphoma cell lines^15,16^. Unexpectedly, NOTCH1 repressed genes were strongly enriched in antigen-processing and presentation, predominately belonging the family of MHC class II genes (Fig. 3b).

**Figure 3:**
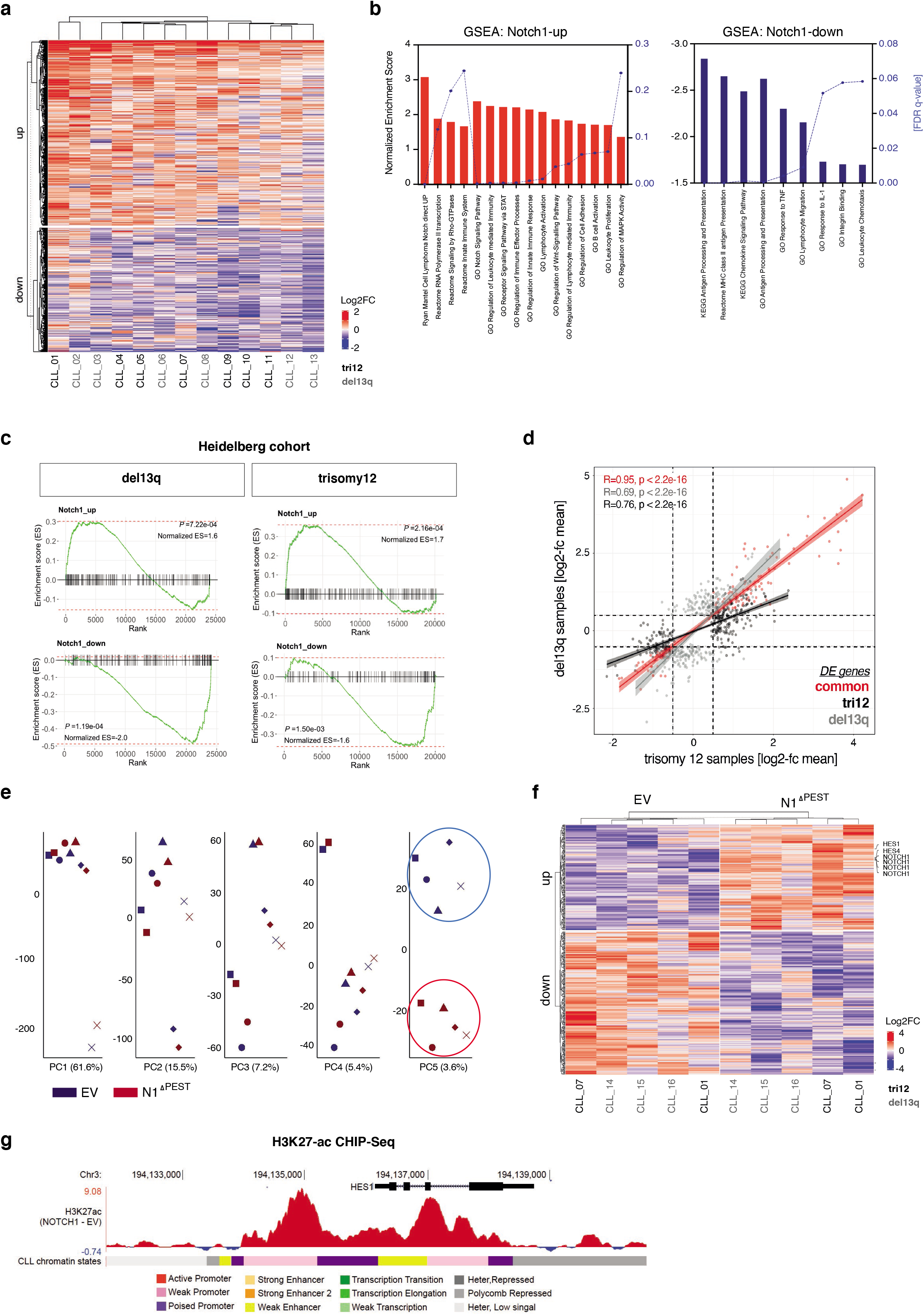
Gene expression profiles of NOTCH1^ΔPEST^ transduced CLL cells. a. Heat map DE genes following NOTCH1^ΔPEST^ overexpression in CLL cells (*n=13*).Libraries were generated from mRNA isolated from FACS sorted GFP^+^ cells 4 days post infection. b. Gene Ontology (GO) and Gene set enrichment analysis (GSEA) results of NOTCH1^ΔPEST^ DE genes identified by RNAseq. Upregulated (left) and down-regulated (right) gene-sets are indicated by bars. Normalized Enrichment Scores are shown on the left black Y-axis, FDR q values on the right Y-axis. c. DE genes identified by RNAseq were validated on a cohort of *NOTCH1*-mutated patients with del13q (left panel) or trisomy 12 (right panel). d. Scatter plot showing the Log2FC mean of DE genes in tri12 (*n=7*, light grey dots, x-axis) vs del13q (*n=6*, black dots y-axis) following NOTCH1^ΔPEST^ overexpression. Red dots represent commonly DE genes between the two different groups. e. Unsupervised principal component analysis of the H3K27ac Chip-seq profiles of five paired CLL primary cases transfected with *expressing vector* or *NOTCH1^ΔPEST^*. 43,300 independent genomic regions were analyzed to generate the PCAs. f. Heatmap shows VST normalized H3K27ac signals for those peaks identified in principal component 5, associated with promoters or gene bodies. g. Example targeting *HES1* gene, identified to have an increase H3K27ac signal in NOTCH1ΔPest expressing samples. The mean value of the subtracted signal calculated for each individual paired sample per 1bp is shown.

While *NOTCH1* mutations have been identified in less than 15% of treatment naïve, unselected patients^1,19^, this frequency is significantly higher in patients with trisomy12, in which *NOTCH1* mutations can be found in 40-50% of patients^26,27^. Importantly, the presence of *NOTCH1* mutations in tri12-CLL is associated with high rates of transformation into Richter’s syndrome^11,28^. The underlying reasons for this peculiar association are unknown, but it suggests that NOTCH1 may regulate distinct, transformation-favoring genes in cells carrying an extra chromosome 12. To address this hypothesis, we separately analyzed the gene expression profiles induced by NOTCH1 in 7 tri12 and 6 del13q patients. Pairwise analysis (applying a cut-off of Log2FC>0.5 in at least half of all samples and of Log2FC>0 in all samples) identified 410 NOTCH1-regulated genes in tri12 (268 up-regulated/142 down-regulated) and 418 genes in del13q (236 up-regulated/182 down-regulated) patient samples. DE genes in each group were then used to validate the NOTCH1-transcriptome on a cohort of *NOTCH1*-mutated tri12 and del13q patients^29^ and showed a significant enrichment in the respected genotype (Fig. 3c). The comparison of NOTCH1-induced genes did not identify a distinct expression profile in tri12 compared to del13q cells, as shown by the correlation of the respective logFCs in both sets of deregulated genes (Fig. 3d). Interestingly, the slope of the tri12-specific genes was clearly smaller than that of del13q-specific genes, suggesting that NOTCH1^ΔPEST^ induced a higher amplitude of gene activation/deactivation in the tri12 genetic context (Fig. 3d, black vs grey line). Furthermore, this analysis recognized 130 genes that were de-regulated at similar levels by NOTCH1^ΔPEST^ in all 13 samples (Fig. 3d, red line). Subjecting this gene-set to GSEA identified transcriptional changes in gene clusters involved in signaling, stress-response and antigen-presentation (Supplementary Fig. 2).

In addition to a clear transcriptional modulation, we next assessed whether NOTCH1^ΔPEST^ was also able to induce epigenetic programming at the level of chromatin regulation. For this, we performed ChIP-seq for histone H3 lysine 27 acetylation (H3K27ac), a bona fide mark for active regulatory elements^30^, in five paired CLL samples (expressing either *EV*-control or *NOTCH1^ΔPEST^*). Unsupervised principal component analysis revealed that the first components of the chromatin activation variability were patient specific. However, the fifth component, which explains 3.6% of the total variability, remarkably separated controls from NOTCH1 ^ΔPEST^ expressing samples, regardless of their genetic background (Fig. 3e). This NOTCH1 ^ΔPEST^-associated signature was composed of 587 H3K27ac peaks. Of them, 422 peaks were located at active chromatin states of CLL reference epigenome samples^31^ and at the promoter or gene body of an annotated gene (see materials and methods). Furthermore, at those differentially acetylated peaks we observed a consistent increase or decrease in H3K27ac signal in all cases with NOTCH1^ΔPEST^ regardless of the cytogenetic background (Fig. 3f). Consistently with the RNAseq data, we identified chromatin activation at several NOTCH1 target genes such as *HES1* (Fig. 3g). These data demonstrate that NOTCH1 mutations not only drive transcriptional changes but also induce an aberrant epigenetic programing of CLL cells.

Collectively, these results indicate that NOTCH1 activation positively regulates gene expression important for B cell activation while simultaneously repressing genes required for antigen-presentation. These effects were not qualitatively different in tri12 cells, but here NOTCH1 effects seemed to be more enhanced compared to del13q cells.

### NOTCH1 represses MHC class II genes via down-regulation of *CIITA*

Our RNAseq analyses indicated that NOTCH1 is associated with reduced expression of genes important for antigen-presentation, including *HLA-DM*, -*DR*, -*DP* and -*DQ*, suggesting silencing of the MHC class II locus on chromosome 6. Indeed, H3K27ac CHIP-seq analysis confirmed that this gene repression was due to epigenetic silencing of the entire HLA-locus (Fig. 4a) and demonstrated that, besides gene activating functions, NOTCH1 can also induce repressive effects on transcription. Assessment of surface HLA-DR expression on an additional 12 primary CLL samples confirmed that NOTCH1 is consistently associated with down-regulated HLA-class II genes (Fig. 4b). To provide further evidence for the gene-repressive functions of NOTCH1, we cultured CLL cells from 4 donors with endogenous exon34 mutations of *NOTCH1* under identical conditions on MM-1-stroma cells in the presence or absence of γ-secretase inhibitors (GSI) to block ligand-mediated activation of NOTCH1. As shown by us and others, stroma cells express NOTCH-ligands^32,33^, which can trigger activation of the NOTCH-pathway. Blockage of NOTCH-activation by GSI treatment induced a significant up-regulation of HLA-DR in *NOTCH1*-mutated CLL, supporting that it can repress the expression of HLA-genes (Fig. 4c). Similar to CLL, *NOTCH1*-mutations have been described in 5-10% of MCL patients, causing expression of a truncated, PEST-domain deleted NOTCH1 protein in the majority of cases and being associated with shorter survival rates^34^. To investigate whether NOTCH1 had similar effects on the expression of MHC class II genes in primary MCL cells, malignant B cells from 4 patients were transduced with *NOTCH1^ΔPEST^* or *EV*- control. Similar to CLL, NOTCH1 down-regulated HLA-DR expression in primary MCL cells (Fig. 4d).

**Figure 4:**
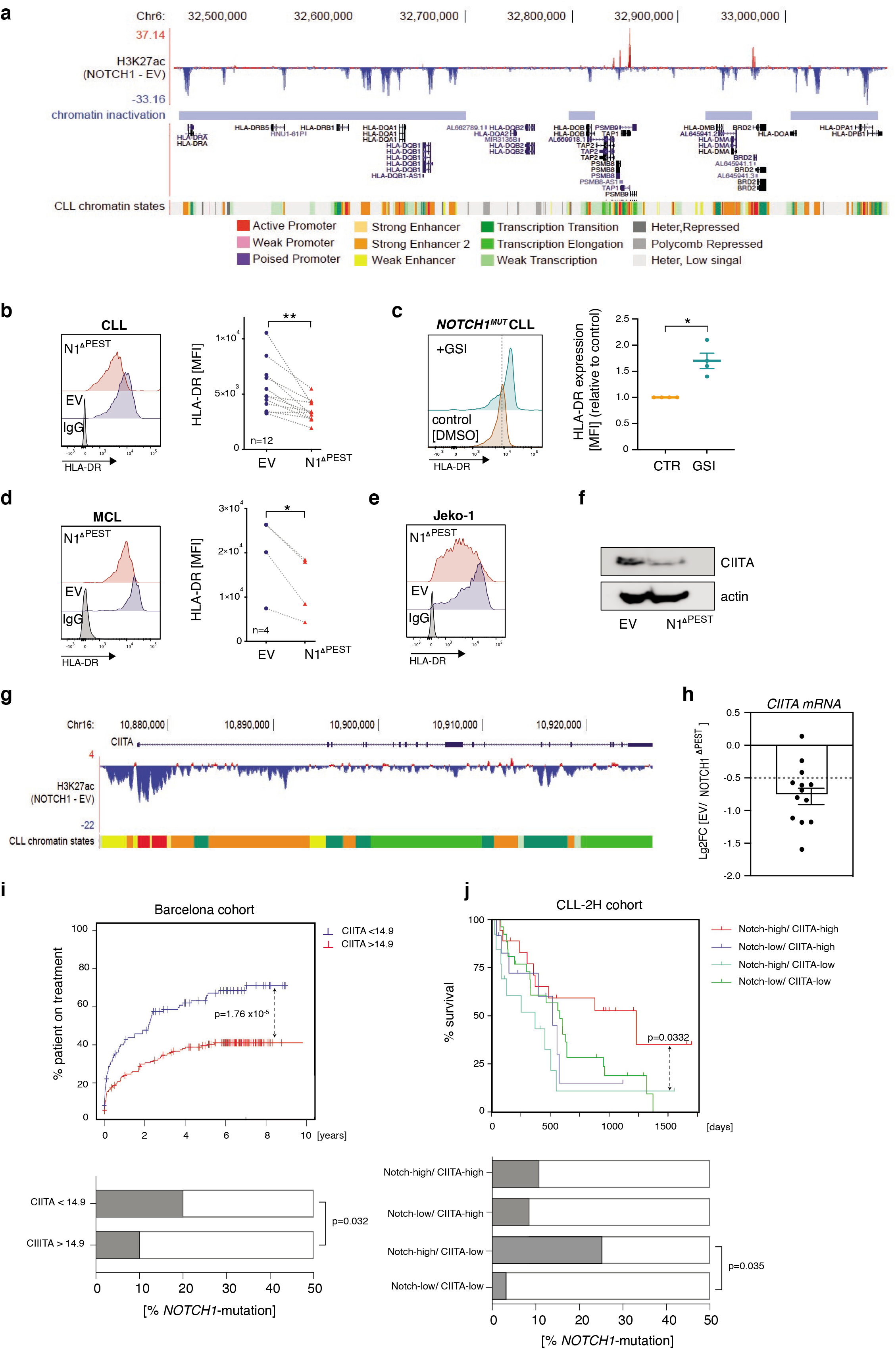
NOTCH1^ΔPEST^ represses MHC class II expression in CLL and MCL. a. H3K27ac Chip-seq profile of the entire MHC class II locus following NOTCH1 ^ΔPEST^ overexpression. The peaks represent the mean of the ratio of values obtained from CLL cells transduced with *NOTCH1^ΔPEST^* or with an *empty vector* control (*n=5*). Each gene and loci locations are shown. b. HLA-DR expression quantified by flow cytometry of transduced CLL cells (*n=12*). HLA-DR expression was analyzed on GFP-positive cells 5 days post transduction. Representative flow cytometry histogram image is shown on the left. c. Quantification of HLA-DR expression on *NOTCH1*-mutated CLL cells treated for 48h with a γ-secretase inhibitor (10μM) or DMSO as control. Relative expression compared to empty vector control is shown (*n=4*). d. HLA-DR expression quantified by flow cytometry of transduced MCL cells (*n=4*). HLA-DR expression was analyzed on GFP-positive cells 5 days post transduction. Representative flow cytometry histogram image is shown on the left. e. Flow cytometry analysis of HLA-DR expression on Jeko-1 cells following overexpression of NOTCH1 ^ΔPEST^. One representative experiment out of three is shown. f. CIITA immunoblot of Jeko-1 cells 5 days post-transduction with *NOTCH1^ΔPEST^* or an *empty vector* control. Proteins were extracted following FACS-of GFP-positive cells. One of three independent experiments is shown. g. H3K27ac Chip-seq profile at *CIITA* locus. The peaks represent the mean of the ratio of values obtained from CLL cells transduced with NOTCH1 ^ΔPEST^ compared to an empty vector control (*n=5*). Chromatin states are color-coded, corresponding to the legend in panel A. h. CIITA mRNA expression analyzed by RNAseq of 13 CLL patients. i. Time to treatment (TTT) curve of a cohort of patients classified as expressing high (red line, CIITA>14.9, *n=156*) or low (blue line<14.9, n=110) CIITA level (upper panel) along with the relative percentage of distribution of NOTCH1 mutated patients in the two subgroups (lower panel). j. Survival of a cohort of patients with a high NOTCH1 gene signature identified by high expression of (*HES1/2, HEY1/2*) and further classified by high or low CIITA expression (upper panel). Relative percentage of distribution of NOTCH1 mutated patients in the 4 sub-groups is indicated in the lower panel. *P < 0.05, **P < 0.01, ***P < 0.001, ****P < 0.0001and not significant (ns) P > 0.05.

The ubiquitous repression of the entire HLA-class II locus suggested that NOTCH1-activity could affect the expression of the class II transactivator (CIITA) in CLL and MCL cells, which is a master regulator for HLA class II genes. CIITA, which itself does not bind to DNA, interacts with multiple transactivating proteins of the MHC class II enhanceosome and regulates gene expression through multiple mechanisms, including recruitment of transcription factor IID, phosphorylation of RNA polymerase II and histone modification (reviewed in^35^). To test whether NOTCH1 affected the protein expression of CIITA, which is also tightly post-transcriptionally regulated, we expressed *NOTCH1^ΔPEST^* in B cell lymphoma cell lines. Importantly, HLA-DR expression was unaffected by NOTCH1 activity in the CLL cell lines MEC-1 and Hg-3 (Supplementary Fig. 3a), further underscoring the limitations of cell lines for studying oncogenic signaling in B cell lymphoma. However, HLA-DR was significantly down-regulated in the MCL cell line Jeko-1, which correspondingly showed a NOTCH1-dependent decreased expression of CIITA protein (Fig. 4e,f), indicating that repression of HLA-class II genes in primary malignant B cells occurred via down-regulation of CIITA. Furthermore, H3K27ac-CHIPseq showed silencing also of the *CIITA* locus on chromosome 16 in all samples (Fig. 4g). Consistently, *CIITA*-RNA levels were significantly down-regulated in primary CLL cells transduced with *NOTCH1^ΔPEST^* (Fig. 4h). In conclusion, our data provide evidence that NOTCH1 down-regulates HLA-class II genes via transcriptional suppression of CIITA.

The downregulation of MHC class II genes provides an immune-escape for cancer cells by reducing their immunogenicity. The clinical significance of the down-regulated or absent HLA-class II expression in B cell lymphoma is illustrated by its negative impact on the prognosis of patients with DLBCL and primary mediastinal B cell lymphoma (PMBCL)^36,37^. To test whether CIITA-dependent down-regulation of MHC class II genes was prognostically important also for CLL, we analyzed whether *CIITA* RNA levels predicted the time to first treatment in a cohort of 266 treatment naïve patients from the International Cancer Genome Consortium (ICGC)^2,38^. Dividing patients into either *CIITA* low or high expresser, based on the overall *CIITA* mRNA abundance, we discovered that those patients with low *CIITA* levels had a significantly more active disease and required treatment sooner. Importantly, *NOTCH1* exon 34 mutations were twice as common in the *CIITA* low expresser group compared to high expresser (Fisher t-test, p> 0.032) (Fig. 4i).

In addition, we assessed the significance of *CIITA* expression in a cohort of pre-treated CLL patients from the CLL-2H study^39^. To also consider NOTCH1-activation in the absence of exon 34 mutations^15^, we first stratified 337 treatment-naïve patients based on the expression of canonical target genes *HES1/2, HEY1/2* and *CIITA* expression (median high vs. median low). This analysis identified a group of patients with high and low expression of *CIITA* in both NOTCH1-activated and non-activated groups, defining 4 patient cohorts (Supplementary Fig. 3b,c). We applied these expression thresholds to gene expression data generated from PBMCs in a cohort of fludarabine-resistant CLL treated in the CLL-2H study. These analyses indicated that high expression of canonical NOTCH-target genes was not per se associated with an unfavorable prognosis, but significantly impacted in the overall survival in combination with low levels of *CIITA* expression (Fig. 4j). Importantly, within the *CIITA^low^* expresser, *NOTCH1* mutations were significantly more frequent in the *NOTCH^high^* versus *NOTCH^low^* group.

Collectively, these data demonstrate that low levels of *CIITA* are associated with a more aggressive disease, in particular for patients with activated NOTCH-signaling. Furthermore, this analysis also suggests that NOTCH1 can be activated in the absence of *NOTCH1*-mutations, as previously reported^15^.

### NOTCH1 up-regulates PD-L1 and impairs T cell activation

NOTCH1-dependent suppression of *CIITA* and further downstream HLA-class II genes indicated a mechanism for an escape from immune surveillance. To provide further evidence for this, we co-cultured primary CLL cells, either transfected with *EV* or *NOTCH1^ΔPEST^*, with Jurkat T cells, expressing luciferase under the control of the NFAT responsive element. Under these conditions, T cell activation was strictly dependent on the presence of the anti-CD19/ anti-CD3 bi-specific antibody Blinatumomab. Notably, expression of *NOTCH1^ΔPEST^* in primary CLL cells mitigated the activation of Jurkat T cells (Fig. 5a). While these results indicated that *NOTCH1* mutations permit immune escape of CLL cells, they also suggested that mechanisms other than HLA-class II down-regulation contribute. To identify potential candidates capable of downregulating TCR activity following NOTCH1-activation we performed quantitative proteomic analysis on primary CLL cells transduced with *NOTCH1^ΔPEST^* or *EV*-control. For the simultaneous analysis of both groups of samples, we applied Tandem Mass Tags (TMT-6plex), which allows for the quantitative comparison between replicates and conditions. Total proteomic investigation of 3 patients recognized 7876 unique proteins and as a result of pairwise analysis, we identified 385 differentially regulated proteins (average Log2FC>0.5 and all 3 patients with Log2FC>0; Fig. 5b). Proteins expressed at a higher level included positive controls such as NOTCH1 and CD38. Interestingly, NOTCH1^ΔPEST^ increased the expression of CD27 and CD274 (PD-L1) in CLL cells, which both can impair T cell activation. Assessment of PD-L1 expression on additional 20 CLL patients, transfected with *NOTCH1^ΔPEST^* or *EV*, invariably showed an up-regulation of PD-L1 through activated NOTCH1. Similar to CLL, primary MCL cells also showed a trend towards an increased expression of PD-L1 following transfection with *NOTCH1^ΔPEST^* (Fig. 5c). Notably, we were unable to recapitulate this phenotype in the CLL cell lines MEC-1 and Hg-3 (Supplementary Fig. 3d), further emphasizing the limitations inherent to studies with cell lines. To demonstrate that NOTCH1^ΔPEST^ functions similarly to ligand-activated, mutated NOTCH1, we performed a reverse experiment by treating *NOTCH1*-mutated CLL with γ-secretase inhibitors (GSI) to block NOTCH-activation. GSI-treatment caused a significant down-regulation of PD-L1 in primary *NOTCH1*-mutated CLL (Fig. 5d).

**Figure 5:**
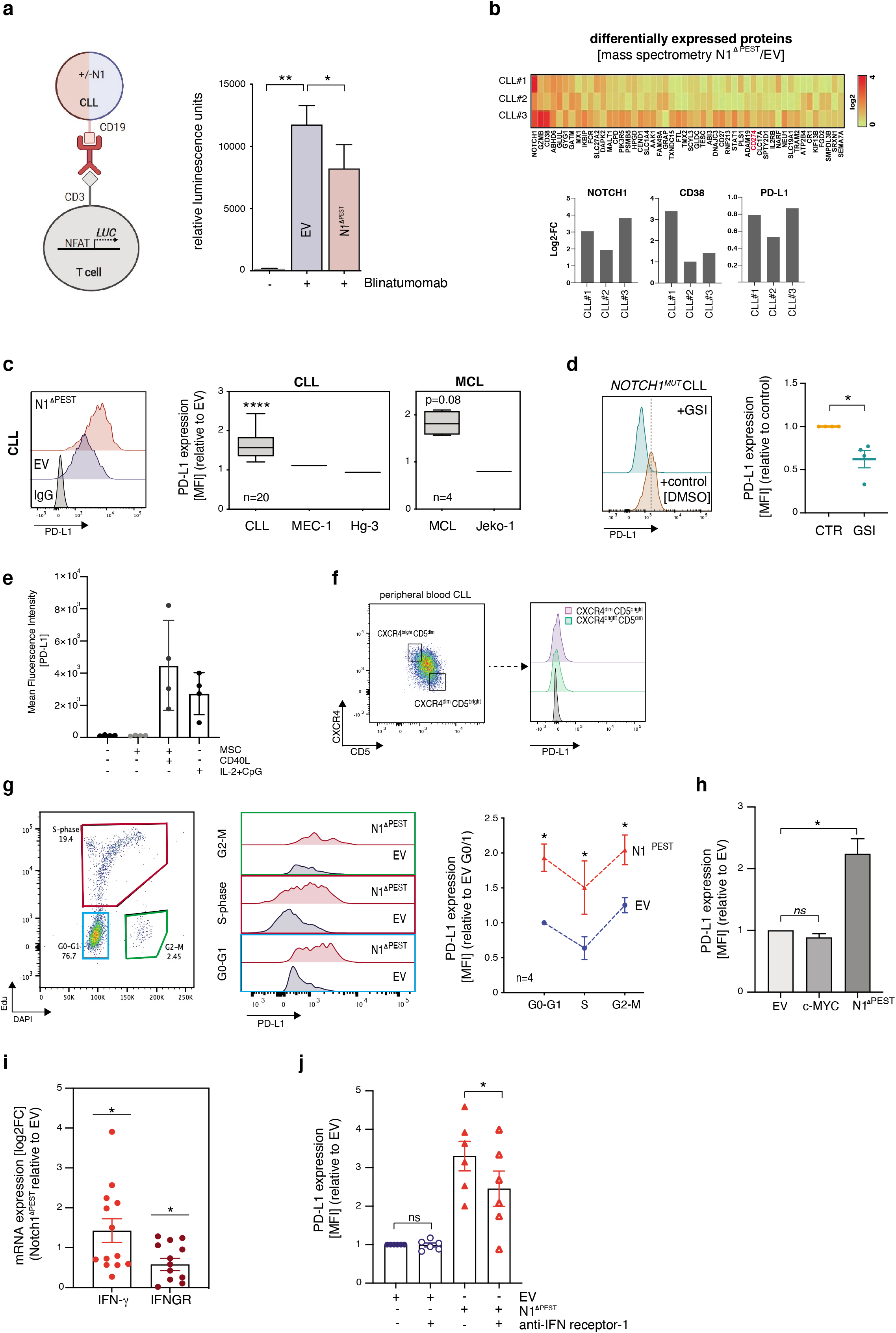
NOTCH1^ΔPEST^ induces the up-regulation of PD-L1 on CLL and MCL cells. a. Quantification of luciferase activity (RLU) of Jurkat NFAT-Luc reporter cells co-cultured with empty vector (blue bar) or NOTCH1 ^ΔPEST^ (red bar) CLL cells in the presence of 10nM Blinatumomab. CLL cells were co-cultured with Jurkat cells at a ratio of 2:5 for 24h. b. Heat map of the top 50 differentially expressed proteins identified by mass spectrometry of NOTCH1^ΔPEST^ transduced cells. Histogram with the Log2FC values from three independent patients analyzed is shown for NOTCH1, CD38 and PD-L1. c. PD-L1 expression quantified as ratio of MFI of CLL (*n=20*) and MCL primary cells (*n=4*)and cell lines transduced with NOTCH1 ^ΔPEST^ or an empty vector control. PD-L1 expression was analyzed on GFP-positive cells 5 days post transduction. Representative flow cytometry histogram images are shown on the left. d. Quantification of PD-L1 on NOTCH1 mutated CLL cells treated for 48h with a γ-secretase inhibitor (10μM) or DMSO as control (*n=4*). Representative flow cytometry histogram image is shown on the left. e. Quantification of PD-L1 expression on activated CLL cells cultured in suspension or on mesenchymal stroma cells for 48h. Alternatively, cells were stimulated with CpGs (ODN-DSP30 1μM) and IL-2 (100 U/ml) (*n=4*) to induce proliferation. f. PD-L1 analysis on freshly isolated CLL cells in CD19^+^CXCR4^dim^/CD5^bright^ and CD19^+^CXCR4^bright^/CD5^dim^ populations (*n=6*). Representative data are shown for one patient and illustrate the gating-strategy and expression of PD-L1. g. PD-L1 expression quantified as ratio of MFI for CLL cell transduced with an *empty vector* (blue) or *NOTCH1^ΔPEST^* (red) during cell cycle progression (*n=4*). Representative flow cytometry and histogram images of each individual cell cycle phase are shown on the left. Cells were pulsed for 12h with Edu prior to fixation. h. Comparison of PD-L1 expression on CLL cells, transduced with *NOTCH1^ΔPEST^* or *c-MYC* relative to the empty vector control (*n=4*). i. Bar graph of the Log2FC values of *IFNG* and *IFNGR1* expression analyzed by RNAseq following NOTCH1^ΔPEST^ transduction (*n=13*). j. PD-L1 expression of CLL cells transduced with an empty vector (blue) or NOTCH1 ^ΔPEST^ (red) treated with a blocking IFNG receptor antibody for 24h. Cohorts are shown as mean ± SD. *P < 0.05, **P < 0.01, ***P < 0.001, ****P < 0.0001 and not significant (ns) P > 0.05.

Importantly, we observed that the constitutive expression of PD-L1 on quiescent cells was minimal, but significantly up-regulated on activated, cycling CLL cells (Fig. 5e), in keeping with a report showing strong expression of PD-L1 on CLL cells in proliferative centers in lymph nodes^40^. To test whether CLL cells recently egressed from lymph nodes had higher expression levels of PD-L1, we assessed its expression on peripheral blood cells expressing CXCR4^dim^/CD5^bright 41^. Unexpectedly, we did not find a different expression of PD-L1 compared to CXCR4^bright^/CD5^dim^ cells, which was overall very low (Fig. 5f). These data demonstrate that the upregulation of PD-L1 in proliferative centers is short-lived.

PD-L1 is also post-transcriptionally regulated through cyclinD-Cdk4 activity, causing cell-cycle dependent oscillations of PDL-L1 expression with a peak expression during M- and early G1-phase^42^. Since NOTCH1^ΔPEST^ provided a significant proliferative advantage for primary CLL cells (Fig. 2c,d), we hypothesized that the increased expression of PD-L1 could be attributed to an increased proliferation, rather than being a specific NOTCH1-response. To address this, we assessed the expression of PD-L1 on cycling CLL cells. For this, cells were stained with an anti-PD-L1 antibody, followed by fixation and staining with Edu to distinguish cells in G0/1 from those in S- or G2/M-phase. In keeping with cell-cycle modulated expression of PD-L1, we observed a significant down-regulation on CLL cells going through S-phase, which recovered with entry into G2/M-phase. Notably, the expression of PD-L1 was consistently increased in *NOTCH1^ΔPEST^*-transduced CLL cells compared to *EV*- controls (Fig. 5g), indicating that the NOTCH1-induced expression of PD-L1 is not dependent on cell proliferation. In addition, although retroviral expression of *c-MYC* caused rigorous proliferation of primary CLL cells (Fig. 1g), c-MYC did not affect PD-L1 expression (Fig. 5h), providing further evidence for a cell-cycle independent regulation of PD-L1 by NOTCH1.

Besides the post-transcriptional regulation of PD-L1, its expression is induced by a variety of pro-inflammatory cytokines, of which interferons are strong inducers. Importantly, CLL cells can produce and secrete IFN-γ, which provides an anti-apoptotic signal through autocrine stimulation ^43^. To investigate whether this feed-back loop was affected by NOTCH1-activation, we analyzed the expression of IFN-γ and its receptor in *NOTCH1^ΔPEST^* transduced cells.

NOTCH1 not only up-regulated *IFN-γ* mRNA, but also the IFN-γ receptor (Fig. 5i), suggesting a contribution of autocrine secreted IFN-γ to NOTCH1-mediated expression of PD-L1. Indeed, CLL cells cultured in the presence of an antibody blocking the IFN-γ receptor showed a significantly down-regulation of PD-L1 (Fig. 5j), which was still significantly higher than PD-L1 levels of *EV*-control cells, indicating that this pathway only partly contributed to the NOTCH1-mediated up-regulation of PD-L1.

In conclusion, these results demonstrate that NOTCH1 signaling promotes escape from immune-surveillance through transcriptional regulation of HLA-class II genes and PD-L1.

### NOTCH1-activation in CLL cells favors expansion of CD4^+^ cells *in vivo*

Our data demonstrated that PD-L1 expression was significantly up-regulated in cycling CLL cells, further enhanced through NOTCH1-activation. To provide *in vivo* evidence supporting this finding, unselected lymph node specimens from CLL/SLL patients were retrieved from the files of the Institute of Pathology, Würzburg, Germany and stained for NOTCH1. We found nuclear expression of NOTCH1 in 12% of all samples (Fig. 6a). Although genomic data for these samples were not available, this frequency is expected based on the occurrence of *NOTCH1* mutations in an unselected patient cohort^19^. For multiparameter analysis of the lymph node microenvironment, we employed imaging mass cytometry (IMC) and applied a panel of isotype-labeled antibodies against B- and T cell epitopes on paraffin-embedded tissues (Fig. 6b). Following cell segmentation using the CellProfiler software we identified areas with high Ki67 signal using HistoCat software^44^ to specifically gate on proliferative centers (PCs). IMC-single cell data identified a high percentage of T cells present in PCs (Fig. 6c). Analysis of PD-L1 expression on CD19^+^ cells revealed a stronger signal in PC areas compared to non-PC areas in all samples, in agreement with published data^40^. Further analyses were restricted to B cells in PCs and showed that nuclear NOTCH1 expression was associated with higher PD-L1 and Ki67 signals, compared to NOTCH1 negative samples (Fig. 6d). Assessment of T cells in PCs also showed a significantly higher infiltration of CD4^+^ cells in NOTCH1-positive samples, associated with higher expression of Ki67, suggesting that NOTCH1-expression in CLL cells promotes T cell expansion. Notably, PD-1 levels on CD4^+^cells were similar between NOTCH-positive and negative samples (Fig. 6e). Similar to CD4^+^cells, CD8^+^ cells were also more abundant in PC of NOTCH1-positive patient samples and they also expressed higher levels of PD-1 (Fig. 6f) in contrast to CD4^+^ cells. These results confirmed our *in vitro* data of NOTCH1-mediated regulation of PD-L1 and indicated that NOTCH1 supports proliferation of CD4^+^ and CD8^+^ cells, with the latter having a more exhausted phenotype.

**Figure 6:**
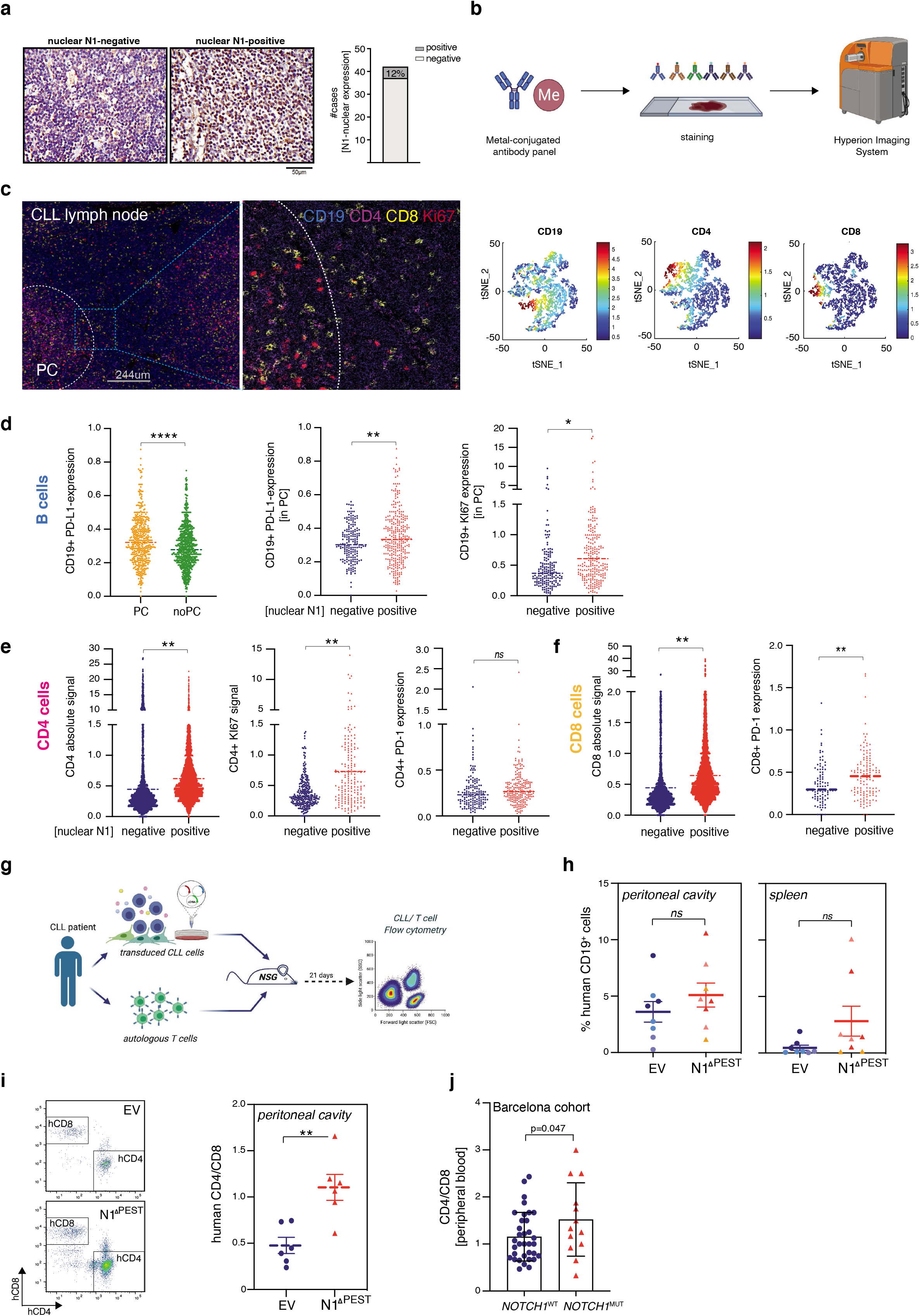
NOTCH1-activation is associated with T cell proliferation *in vivo*. a. IHC staining of NOTCH1 in CLL lymph node biopsies. Five out of 42 samples showed nuclear expression of NOTCH1, representing 12% of cases from an unselected cohort (bar graph). Representative IHC images of positive and negative NOTCH1 nuclear staining are shown. b. Graphical scheme of the procedure used for the mass cytometry analysis. In brief, samples negative (*n=3*) or positive (*n=4*) for nuclear NOTCH1 IHC staining were processed and stained with a panel of metal-tagged antibodies. Following Hyperion Imaging System analysis, the data was processed using Cell Profiler and Histocat software. c. Multiplexed IMC image example of a NOTCH1 positive specimen with an enlarged area of the proliferative center (PC). Scale bar = 244μm. Representative t-SNE plots generated with Histocat showing CD19, CD4 and CD8 cell populations are displayed on the right. d. Intensity of cellular signal per given cell was calculated using the HistoCat software. PD-L1 signal in CD19 gated cells following tNSE analysis in PC and non-PC area (i), PD-L1 (ii) and KI67 (iii) in CD19-gated cells following tNSE analysis in PC of samples with a positive or negative NOTCH1 nuclear staining. e. Total CD4 signal in PC (i) and KI67 (ii) and PD-1 (iii) in CD4-gated cells following tNSE analysis in PCs of NOTCH1 positive or negative samples. f. Total CD8 signal in PC (i) and PD-1 (ii) in CD8-gated cells following tNSE analysis in PCs of NOTCH1 positive or negative samples. g. Graphical scheme of the *in vivo* experiment using transduced CLL cells and autologous T cell to generate NSG chimera. CLL cells were transduced with an *empty vector* or *NOTCH1^ΔPEST^* and intraperitoneally injected into NSG mice. Autologous T cells were cultured with IL-2, a-CD3 and a-CD28 for 7 days prior to injection. h. Engraftment of human CD19^+^ cells in the peritoneal cavity (left) and spleen (right) of mice injected with NOTCH1^ΔPEST^ (n=8) or an empty vector (n=8) transduced CLL cells. In total 16 mice were analyzed using cells from 3 independent donors. i. Quantification of the ratio of human autologous CD4^+^/CD8^+^ T cells in the peritoneal cavity of mice injected with CLL cells with or without NOTCH1 ^ΔPEST^. Mean value was obtained from 3 independent experiment. In total analysis was performed on 6 mice injected with 3 independent CLL donor cells. j. Ratio of human CD4^+^/CD8^+^ T cells in the peripheral blood of a cohort of naive CLL patients with NOTCH1 wild type or mutated Cohorts are shown as median (D-E) and as mean ± SEM (H-J). *P <0.05, **P < 0.01, ***P < 0.001, ****P < 0.0001 and not significant (ns) P > 0.05.

To provide further experimental evidence for this hypothesis, we injected *NOD.Cg-Prkcd ^scid^Il2rd^tm1Wjl^/ Szj* (NSG) mice intraperitoneally with isogeneic primary CLL cells, carrying either *NOTCH1^ΔPEST^* or *EV*-control. Prior to this, autologous T cells were isolated with anti-CD3 beads, cultured for 7 days and then co-injected at a ratio of 20:1 (CLL:T cell) (Fig. 6g). Assessment of engrafted CLL cells showed a moderate, but not significant, increase in the tumor burden of NOTCH1-transduced cells after 3 weeks (Fig. 6h). Importantly, we observed a significant increase of CD4^+^ T cell in the peritoneal cavity of mice diseased with *NOTCH1^ΔPEST^*-expressing CLL cells, demonstrating that NOTCH1 promotes expansion of CD4^+^ cells (Fig. 6i). This is further supported through clinical data from untreated CLL patients, showing a higher ratio of CD4^+^/CD8^+^ cells in the peripheral blood of *NOTCH1* mutated patients compared to NOTCH-wild type patients (Fig. 6j).

## Discussion

Numerous sequencing studies have identified many mutations recurrently found in malignant B cells from CLL and MCL-patients^1,19,34^. To translate this knowledge into patient care, functional studies are needed to understand the mechanisms governed by these mutations and to identify downstream effects amenable for therapeutic interventions. Here we provide a method to functionally interrogate gene-mutations in primary human malignant B cells. For a disease such as MCL or for studying tumor cells with structural chromosomal abnormalities, for which no animal models exist, this method is indeed a unique opportunity to decipher the underlying disease biology.

We applied this technique to address the question why CLL and MCL patients carrying *NOTCH1* mutations have a dismal prognosis. Previous studies had approached this question through investigations in cell lines, commonly derived from therapy-resistant patients. Undoubtably, these studies have made significant contributions to our understanding of the NOTCH1 biology and described that it promotes proliferation, BCR-signaling, MAPK-signaling and chemotaxis in CLL cells^15–17,21,45^. Our studies with primary malignant B cells indeed confirmed these findings, but also identified yet unappreciated roles of NOTCH1.

Since the majority of *NOTCH1* mutations in CLL and MCL do not affect protein binding to DNA but instead impair its proteasomal degradation by truncating the PEST domain^2,19,34,46^, several aspects need to be considered to fully comprehend when, where and how NOTCH1 signaling drives disease progression. The preserved DNA-binding function of mutated NOTCH1 suggests that it regulates the expression of identical genes than wild-type NOTCH1 and that disease-promoting events are rather caused through secondary events attributed to signal-persistence. Thus, mutated NOTCH1 still requires binding of NOTCH-ligands, expressed in trans, for signaling. This apparent ordinary ligand-receptor signaling is complicated by an unusual redundancy of ligands, expressed not only on cells in the tumor microenvironment, but also on malignant B cells themselves^33^, as well as co-expression of other NOTCH-receptors on tumor B cells^20^. The simultaneous expression of both NOTCH receptors and ligands on the same cell is known to lead to cis-inhibition^47^. NOTCH-signal strength is therefore dependent on the balance between cis-inhibition and trans-activation. Unfortunately, in CLL as well as in MCL, very limited knowledge exists about the expression of NOTCH-ligands and receptors in distinct niches *in vivo*. Extrapolating from our recent *in vitro* data, ligand and receptor expression are also likely to be dynamic *in vivo* and regulated by NOTCH-signaling itself^33^. Another unknown variable, essential for understanding how *NOTCH1* mutations modulate disease biology is the length of time a tumor cell resides in one tissue before migrating to another, which will almost certainly impact on the activation of NOTCH1 in tumor cells as signaling is cell-contact dependent.

We believe this complexity of NOTCH1-signalling is important for understanding the recently reported activation of NOTCH1 in 50% of CLL patients, based on the presence of NOTCH1-ICD protein, although PEST-truncating mutations of *NOTCH1* were only found in 22% in the same study^15^. While the reasons for the discrepancy between the presence of NOTCH1-ICD protein and gene-mutations remains elusive, these data also indicate that carrying a *NOTCH1* mutation is fundamentally different from expressing NOTCH1-ICD as only gene mutations appear to predict for a poor clinical outcome. Furthermore, these data also hint on the importance of the tumor microenvironment for NOTCH1-activation, suggesting that mutations of oncogenes still rely on signals from non-malignant cells to fully unfold their detrimental effects. Our limited knowledge about these factors contributing to the activation of mutated NOTCH1 in CLL suggest that the lymph node environment is predominantly important for its activation^17,22^.

The negative prognostic effect of *NOTCH1* mutations in CLL becomes even more evident if they occur on the background of trisomy 12. NOTCH1 mutations are enriched in patients with trisomy 12 ^26,27^, suggesting that this chromosomal aberration provides a selective advantage for *NOTCH1* mutations. In addition, the risk for patients with trisomy 12 for transformation into clonally related Richter’s syndrome is 10-times higher for NOTCH1-mutated patients compared to wild-type, suggesting that NOTCH1-signaling drives genomic instability and clonal evolution^11,48^. Our method to generate isogenic primary CLL cells provided an opportunity to directly address this question. Unexpectedly, our experiments did not identify a distinct NOTCH1-regulated gene-set, present only in trisomy 12 cells, but rather indicated a higher amplitude of gene-regulation in trisomy 12 compared to del13q cells. This observation raises a further question of what other factors determine the selective advantage for clones concurrently harboring trisomy12 and NOTCH1-mutations? A possible explanation is the observation that trisomy 12 CLL cells have an increased expression of CD29, CD49d and ITGB7, which occurs independently of NOTCH1 mutations^49^ and allows for an improved adherence to cells of the microenvironment. As an immediate consequence and since *NOTCH1*-mutated cells are still dependent on ligand-binding for activation, trisomy 12 cells may experience prolonged NOTCH1-signaling. Therefore, and based on our data, we propose that in the subgroup of patients with trisomy 12 the selective advantage for *NOTCH*-mutations is based on enhanced, ligand-mediated NOTCH1-activation, rather than due to a specific genetic program governed by NOTCH1.

Gene repressive functions of NOTCH1 have previously been underappreciated. Our data indicated that NOTCH1 signaling permits immune escape of malignant B cells through down-regulation of HLA-class II expression. The prognostic significance of HLA-class II expression is well documented for DLBCL^50^ and PMBCL^37^ and shows that short overall survival is associated with low expression levels in these entities. Similar to our study, low HLA-expression levels were not due to large genetic deletions on chromosome 6 but correlated with *CIITA* expression levels^36^, pointing to transcriptional de-regulation of HLA class II genes in high grade lymphoma.

Our data suggest that *CIITA* expression levels are a novel prognostic marker also for indolent B cell malignancies and show that NOTCH1 is a strong suppressor of *CIITA* transcription. NOTCH1-mediated control of *CIITA* expression has not previously been reported and the underlying mechanisms of this regulation remain to be defined. Since we observed that NOTCH1-signaling also up-regulated IFN-γ, which itself activates *CIITA* transcription^51^, the transcriptional repression of *CIITA* by NOTCH1 likely involves epigenetic silencing of its promotors as shown for other hematological malignancies^52^.

The NOTCH1-mediated modulation of surface receptors regulating the interaction with T cells expectedly has effects on the composition of the tumor microenvironment. We found significantly more cycling CLL cells in proliferative lymph nodes of NOTCH1-expressing cells compared to non-expresser, associated with an increased number of CD4^+^ and CD8^+^ T cells. This association is likely to be mediated through the recruitment of T cells through the secretion of CCL3 and CCL4, derived from activated CLL cells ^53,54^. Notably, the increased number of CD8^+^ T cells, but not of CD4^+^ T cells, was associated with higher expression of PD-1 in NOTCH1 positive samples, indicating terminal differentiation and exhaustion of CD8^+^cells. The role of CD4^+^ T cell-subsets in CLL is far from being completely understood, but the collective evidence indicates that CD4^+^ T cells overall are tumor-promoting. This conclusion is based on the dependency of CLL cells on autologous T cells to engraft in NSG mice^55^, *in vitro* growth promoting effects of CLL-specific Th1-cells^56^ and correlation between higher CD4^+^ cell counts and shorter PFS and OS^57^. The relative contribution of individual CD4^+^ subsets is less clear, but numerous studies suggest that this phenotype may predominantly be driven by Tregs^58,59^, possibly through their secretion of pro-leukemic cytokines such as TGFβ and IL-10. While the comparison of data from human and mouse always needs to be done with caution, data from our experiments in NSG mice indicate that NOTCH1-signaling in CLL cells drive the expansion of CD4^+^ T cells, in keeping with those studies.

In contrast to the CIITA-HLA-class II axis, the role of PD-L1 for the suppression of T cell functions in CLL is better defined. Pre-clinical data indicate that the PD-1/ PD-L1axis actively contributes to immune escape, demonstrating that PD-L1 inhibition prevented the development of a CLL-like disease in the Eμ-TCL1 mouse model^60^. This therapeutic effect was further enhanced through a simultaneously administered BTK-inhibitor^61^. Additionally, PD-1 blockage restored normal immune synapse formation between T and CLL cells^62^. A recent study demonstrated that activation of autologous T cells with an E3-ligase inhibitor also reverted PD-L1 mediated suppression of cytotoxic T cells, causing anti-tumor effects in a CLL xenograft model^63^. Although these pre-clinical data strongly suggest that immune checkpoint blockage is therapeutically useful, clinical data supporting this have been inconclusive to date. A phase II clinical trial with pembrolizumab, a humanized anti-PD-1 antibody, failed to show objective responses in non-transformed CLL. However, the same study reported overall response in 44% of patients with Richter’s transformation (RT)^64^. The reasons for this discrepancy are unknown; enhanced immunogenicity of RT cells may be based on the presentation of tumor-neoantigens, generated in the process of transformation. While the presence of *NOTCH1* mutations is strongly associated with Richter’s transformation, it remains unclear from this trial whether responding patients were carrying *NOTCH1* mutations. Our data predict that PD-1/ PD-L1 inhibition could be more efficacious in *NOTCH1* mutated patients and future prospective studies are needed to address this.

## Acknowledgements

We would like to express our deePEST gratitude to patients who donated blood for research purposes. In particular, we thank Dr Joanna Baxter and her team for enrolling patients into these studies. We also wish to thank the Cambridge NIHR BRC Cell Phenotyping Hub for their advice and support in cell sorting, Dr Richard Grenfell (CRUK Cambridge Institute) for his support with the Hyperion tissue imager and Dr Leigh-Anne McDuffus for her help with IMC processing and analysis. This work was funded by Cancer Research UK (CRUK; C49940/A17480-I.R. is a senior CRUK fellow), Kay Kendall Leukaemia Fund (M.M-KKL1258), and Fundació La Marató de TV3 (201924-30). A.M.D is supported by the Beatriu de Pinós Programme of the Government of Catalonia (2018-BP-00231). This work was partially developed at the Centro Esther Koplowitz (CEK, Barcelona, Spain).

## Author Contributions

M.M performed and analyzed experiments. A.M.D, S.C. and J.I.M.S. ran and analyzed the H3K27ac ChIP-seq., E.G.H. and A.R. performed and analyzed the IHC from CLL lymph nodes. J.B., J.L., S.S. and T.Z. analyzed gene expression in primary CLL samples. A.Mo. helped to perform the PDX experiment. V.N.R.F, C.S.R.C and C.DS. ran and analyzed mass spectrometry experiments. I.M. and S.C. analyzed RNAseq data. S.D. provided conceptional input and human CLL samples. This project was designed by J.I.M.S and I.R., I.R. wrote the manuscript.

## Methods

### Primary cells and cell culture

After patients’ informed consent and in accordance with the Helsinki Declaration, peripheral blood was obtained from patients with a diagnosis of CLL or MCL. Studies were approved by the Cambridgeshire Research Ethics Committee (07/MRE05/44).

PBMCs were isolated from heparinized blood samples from patients by centrifugation over a Ficoll-Hypaque layer (PAN-Biotech, Aidenbach, Germany). Purity of CLL population was assessed by flow cytometry and only samples with >85% CD19^+^CD5^+^ were used. After harvesting, malignant B cells were either frozen down as viable cells or directly cultured in Advanced Roswell Park Memorial Institute medium (Advanced RPMI-1640; Invitrogen, Carlsbad, CA) with GlutaMAX containing 10% FBS (Gibco), 100 IU/ml penicillin and 100 μg/ml streptomycin and kept at 37 °C in a humidified incubator (5% CO_2_ and 95% atmosphere). We have not observed differences in transduction-efficacy between fresh and frozen cells.

Autologous patient derived T cells were isolated using CD3 MicroBeads (Miltenyi Biotec) according to manufacturer instructions. Purified cells were then cultured for 7 days at the density of 10^6^ cells in RPMI supplemented with 10% FBS, 1000 unit/mL IL-2 (R&D systems), anti-CD3 (2ug/mL), anti-CD28 (4ug/mL) 100 IU/ml penicillin and 100 μg/ml streptomycin and kept at 37 °C in a humidified incubator.

MM-1 feeder cells were cultured in MEM Alpha+GlutaMAX medium (ThermoFisher Scientific, Winsford, UK) supplemented with 10% FBS (Gibco), 10% horse serum (Sigma-Aldrich, Dorset, UK), 10 μM 2-ME and 1% penicillin/streptomycin (Gibco).

Cell lines MEC-1, Hg-m3, Jeko-1 and Jurkat were cultured in RPMI-1640 (Invitrogen, Carlsbad, CA) supplemented with 10% fetal calf serum (Gibco), 100 IU/ml penicillin and 100 μg/ml streptomycin (Gibco). Lenti-x-293 Cell Line (Clontech Laboratories, 632180) were cultured in Dulbecco’s modified Eagle’s medium (DMEM, Invitrogen, Carlsbad, CA) containing 10% FBS, 100 IU/ml penicillin and 100 μg/ml streptomycin and kept at 37 °C in a humidified incubator (5% CO_2_ and 95% atmosphere). All cell lines used in this study were tested to be free from mycoplasma.

### Mouse model

8–10-week-old male NOD.*Cg-Prkdc^scid^Il2rg^tm1Wjl^/SzJ* (NSG) mice injected intraperitoneally with 10^7^ retrovirally transduced CLL cells and 5*10^5^ autologous T cells (20:1 CLL:T cell). Following close monitoring for 3 weeks mice were culled and spleen and peritoneal cavity fluid were harvested for the analysis of human cell engraftment.

These animal studies have been regulated under the Animals (Scientific Procedures) Act 1986 Amendment Regulations 2012 following ethical review by the University of Cambridge Animal Welfare and Ethical Review Body (AWERB-PPL number P846C00DB).

### Flow cytometry

Cells were stained with fluorophore-labelled antibodies in 2% BSA in PBS according to the manufacturer’s instructions. For apoptosis analysis, conjugated Annexin-V and DAPI were used for the detection of apoptotic cells according to the manufacturer’s instructions. Cell cycle analysis was performed using the Click-iT™ EdU Alexa Fluor™ 647 Flow Cytometry Assay Kit (ThermoFisher Scientific) according to manufacturer’s instructions. Cells were pulsed with 10μM Edu for 12 hours.

Calcium Flux assay was performed using 5×10^6^ cells. Fluo-4 (5μM) (ThermoFisher Scientific) was added to 500ul of cells in serum free media and incubated for 15 minutes at room temperature with protection from light. Cells were then washed and re-suspended in 100 ul HBSS (Ca2+ free) plus 20ug biotin-SP AffiniPure Fab Fragment Goat Anti-Human IgM for 20 minutes on ice. Cells were then washed, re-suspended in 500ul HBSS and incubated for 20 minutes at 37 °C. DAPI was added to identify dead cells. Samples were analyzed on flow cytometry. Initial measurement was lasting for 20 seconds to record baseline Ca^2+^ signal, then 20ul streptavidin (1mg/ml) was added to stimulate the Ca^2+^ flow. Measurement was resumed for up to 180 seconds.

Samples were acquired on a LSRFortessa™ X-20 cell analyzer (BD Biosciences, Oxford, UK) and analyzed using FlowJo software (Tree Star).

FACS cell sorting was performed using the BD Influx™ Cell Sorter (BD Biosciences).

### T cell reporter assay

T Cell Activation Bioassay (Promega) was performed according to the manufacturer’s instructions. Four days post transduction CLL cells were sorted for GFP^+^ and CD19^+^expression. Following sorting, 2*10^4^ CLL cells where then co-cultured with 5*10^4^ Jurkat NFAT reporter cells with the addition of Blinatumomab (10nM) in white walled 96 wells plate (Corning). TCR-mediated luminescence was measured 24h later using SpectraMax M5e Microplate Reader.

### Expression analysis/qPCR

Total RNA was isolated using the RNeasy Mini Kit (Qiagen, Manchester, UK), and complementary DNA (cDNA) was obtained using the qScriptTM cDNA SuperMix kit (QuantaBio, Beverly, MA, USA). Quantitative reverse-transcription polymerase chain reaction (RT-qPCR) was performed on isolated mRNA using the fast SYBR reagents and the Applied Biosystems™ QuantStudio™ 12K system. Target gene expression levels were normalized to GAPDH and values are represented as fold change relative to control using the ΔΔCt method.

### Western blot

Cultured cells were collected and lysed with RIPA buffer and a total of 20 μg protein was separated by SDS-polyacrylamide gel electrophoresis using 4-12% NuPAGE Bis-Tris gels (ThermoFisher Scientific), blotted to polyvinylidene difluoride (PVDF) membranes (Millipore), and probed with primary antibodies (B-Actin-HRP (Cell Signalling), CIITA 7-1H (Insight Biotech)). Images were captured with the Azure Biosystem c300 (Dublin, CA, USA) digital imaging system.

### Mass spectrometry

Following cell sorting 10^6^ cell pellets were collected for mass spectrometry analysis. Protein isolation and TMT-6plex labelling was performed as described previously^65^. TMT mix was fractionated on a Dionex Ultimate 3000 system at high pH using the X-Bridge C18 column (3.5μm, 2.1x×150mm, Waters) with 90min linear gradient from 5% to 95% acetonitrile containing 20mM ammonium hydroxide at a flow rate of 0.2ml/min. Peptides fractions were collected between 20-55 minutes and were dried with speed vac concentrator. Each fraction was reconstituted in 0.1% formic acid for liquid chromatography tandem mass spectrometry (LC–MS/MS) analysis.

Peptide fractions were analyzed on a Dionex Ultimate 3000 system coupled with the nano-ESI source Fusion Lumos Orbitrap Mass Spectrometer (Thermo Scientific). Peptides were trapped on a 100μm ID X 2 cm microcapillary C18 column (5μm, 100A) followed by 2h elution using 75μm ID X 25 cm C18 RP column (3μm, 100A) at 300nl/min flow rate. In each data collection cycle, one full MS scan (380–1,500 m/z) was acquired in the Orbitrap (120K resolution, automatic gain control (AGC) setting of 3×105 and Maximum Injection Time (MIT) of 100 ms). The subsequent MS2 was conducted with a top speed approach using a 3-s duration. The most abundant ions were selected for fragmentation by collision induced dissociation (CID). CID was performed with a collision energy of 35%, an AGC setting of 1×10^4^, an isolation window of 0.7 Da, a MIT of 50ms. Previously analyzed precursor ions were dynamically excluded for 45s. During the MS3 analyses for TMT quantification, precursor ion selection was based on the previous MS2 scan and isolated using a 2.0Da m/z window. MS2–MS3 was conducted using sequential precursor selection (SPS) methodology with the top10 settings.

For MS3, HCD was used and performed using 65% collision energy and reporter ions were detected using the Orbitrap (50K resolution, an AGC setting of 1×10^5^ and MIT of 105 ms). Peptide intensities were then normalized using median scaling and protein level quantification was obtained by the summation of the normalized peptide intensities. A statistical analysis of differentially regulated proteins was carried out using the limma R-package from Bioconductor ^66^Multiple testing correction of p-values was applied using the Benjamini-Hochberg method (https://www.jstor.org/stable/2346101?seq=1#page_scan_tab_contents) to control the false discovery rate (FDR). The mass spectrometry proteomics data have been deposited to the ProteomeXchange Consortium via the PRIDE ^67^ partner repository with the dataset identifier PXD024112.

### Mass cytometry

FPPE lymph node Sections 5μm in thickness were cut with a Leica CM 1850 UV cryomicrotome and processed according to the manufacturer’s instruction. In brief, slides were baked for 2h at 60 °C before dewaxing with xylene and hydration with descending grades of ethanol. Tissue sections were then incubated with the antigen retrieval solution (pH 9) (Abcam) for 30 minutes, blocked with 3% BSA in PBS for 45 minutes at room temperature and then incubated with Fluidigm pathologist-verified Maxpar antibodies overnight at 4°C in a humidified chamber. The following day, the slides were washed in 0.2% Triton X-100, followed by PBS and then stained with DNA intercalator-Ir (1:2,000 dilution; Fluidigm) for 30 min at room temperature. Slides were washed in distilled deionized water and air-dried for ~30 min. Slides were inserted into the Hyperion Imaging System (Fluidigm) for data acquisition. (https://www.fluidigm.com/binaries/content/documents/fluidigm/resources/imaging-mass-cytometry-staining-for-ffpe-sections-400322-pr/imaging-mass-cytometry-staining-for-ffpe-sections-400322-pr/fluidigm%3Afile)

For the staining we used the following Fluidigm pathologist-verified Maxpar antibodies: Anti-CD19 (6OMP31)-142^Nd^; -Anti-Human CD4 (EPR6855)-156^Gd^; -Anti-Human CD8a (D8A8Y)-162^Dy^; -Anti-Human PD-1 (EPR4877(2))-165^Ho^; -Anti-Ki-67 (B56)-168^Er^; -Anti-Human PD-L1 (E1L3N)-150^Nd^; -Anti-Pan-Actin (D18C11)-175^Lu^; -Anti-Histone 3 (D1H2)-176^Yb^

Images acquired with the Hyperion Imaging System were reviewed and single ROI were exported using MCD Viewer (Fluidigm v1.0.560.6). Single cell segmentation was performed using the open-source software CellProfiler (Broad Institute). For this, individual nuclei were identified using the DNA staining intercalator-Ir and Histone H3 marker followed by identification of the cellular region by a circle of a defined radius. From this, we could now measure the intensity in each channel, and thus a proxy of the expression level of the protein in each individual cell. As the dynamic ranges of the different channels vary considerably our analysis was limited to the comparison of each single channel across the different tissue sections. In order to obtain the intensity of each channel we used the Histology Topography Cytometry Analysis Toolbox (HistoCat) software^44^. Area with high density of KI67 expression were considered proliferation centers. t-SNE analysis across the markers of interest was created and single channel heatmaps were generated in order to gate on specific cell types. Raw data of each population was extracted into excel files and plotted using GraphPad Prism 9.0 (GraphPad Software, La Jolla, USA).

### RNA-seq

Total RNA was isolated from GFP^+^CD19^+^ sorted cells using the RNeasy Mini Kit (Qiagen, Manchester, UK). Samples (25 ng total RNA) were then processed for NGS sequencing using the NuGEN TRIO Kit (NuGEN) and the size distribution of the resulting libraries was analyzed on Agilent Bioanalyzer HS DNA chips. A single library pool containing all samples was generated for sequencing and quantified using the NEB Library Quant kit, a SYBRgreen based qPCR method.

The sequencing of the library pool was performed in two runs on both lanes of a HiSeq 2500 RapidRun flow cell in the paired-end mode: 101 cycles for read 1, 9 cycles for the index read and another 101 cycles for read 2. Both runs generated excellent read qualities and quantities as indicated by the Illumina SAV software tool. Bcl-to-Fastq conversion and de-multiplexing of the reads were performed with the Illumina CASAVA 1.8.2 software using standard settings. For all analyses, quality checks were performed using FastQC (https://www.bioinformatics.babraham.ac.uk/projects/fastqc). The alignment to the reference genome (human genome hg38, genome assembly GRCh38.p13) was done using STAR version 2.5.2a (https://dx.doi.org/10.1093/bioinformatics/bts635). Pre- and post-alignment quality checks were summarised using MultiQC. Gene expression counts were obtained using featureCounts version v1.6.0 (https://doi.org/10.1093/bioinformatics/btt656). Additional quality checks include MA plots and heatmaps representing the Jaccard Similarity Index (JSI).

The normalization of expression levels was performed using quantile normalization using the function normalize.quantiles from the R package preprocessCore (https://github.com/bmbolstad/preprocessCore), followed by edgeR internal normalization. The differential expression analysis was performed using the standard functions from edgeR pipeline, version 3.28.0.

The Fold Change (FC) of the normalized counts of all the genes per pair of samples was calculated as B/A, where B are the NOTCH1 mutated samples and A the control samples.

The differentially expressed (DE) genes were identified as follow: -upregulated those genes that had a log2FC value > 0.5 in at least half of the samples while as down-regulated the ones with a log2FC value < −0.5 in at least half of the samples. A more stringent cut-off was used for groups of samples with tri12 or del13q mutation. As up-regulated genes were characterized those genes that had a log2FC value > 0.5 in at least half of the samples and >0 in all the samples, while as down-regulated the ones with a log2FC value < −0.5 in at least half of the samples and <0 in all the samples. The Pearson correlation coefficients were calculated between the expression of the DE genes of the two groups.

### H3K27ac ChIP-seq

10^6^ cells were FACS sorted 5 days following transduction. The samples were immediately cross-linked in 1% Formaldehyde and the reaction was then quenched with 0.125M Glycine. The fastq files of the ChIP-seq data were aligned to genome build GRCh38 (using bwa 0.7.7, picard and samtools) and wiggle plots were generated (using PhantomPeakQualTools) according to the Blueprint pipeline (http://dcc.blueprint-epigenome.eu/#/md/methods).

Peaks of the H3K27ac data were called as described (http://dcc.blueprint-epigenome.eu/#/md/methods) using MACS2 (version 2.0.10.20131216). For all samples, H3K27ac peaks were called without input control. A set of consensus peaks for all the samples was generated by merging the locations of the separate peaks per sample. Variance Stabilized Transformed (VST) values were calculated for the consensus peaks using DESeq2. For downstream analyses, only peaks present in at least 2 samples were used (45.300 peaks). All Principal Component Analyses (PCAs) were generated with the prcomp function using corrected VST values.

For the isolation of the peaks forming principal component 5 (PC5), the Pearson correlation coefficients between the eigenvalues of PC5 and each peak across all samples were calculated. Correlation coefficients with a p value < 0.05 were included, resulting in 587 peaks. Using previously reported chromatin states of reference CLL samples, peaks located in inactive chromatin regions were removed resulting in 484 peaks. All known genes of the hg38 annotation were downloaded using the GenomicFeatures package, and their locations were extended by 1.5 kb upstream to include their promoter region. The peaks were subsequently annotated according to overlaps with the genes’ coordinates, resulting in 422 peaks. The rtracklayer package was used for the import of the H3K27ac signal files of the samples. The subtracted signal was calculated in each pair of samples (NOTCH1 mutated - control) per 1bp. The bedGraphToBigWig application was used for the transformation of those regions to signal files appropriate for loading to the UCSC browser.

### Patients validation cohort

Data from a published gene expression dataset ^29^ was re-analyzed based on NOTCH1 mutational status and genetic background. Differentially expressed genes between NOTCH1 mutated samples and NOTCH1 wild type samples were identified using DESeq2 (https://genomebiology.biomedcentral.com/articles/10.1186/s13059-014-0550-8). The R package FGSEA (http://bioconductor.org/packages/release/bioc/html/fgsea.html) was then used for gene set enrichment analysis against gene defined as up-regulated and down-regulated after NOTCH1 over-expression by RNAseq.

### Clinical association of CIITA expression

CIITA expression was correlated with genes indicating NOTCH-pathway activation (averaged expression levels of *HES1/2, HEY1/2*) on n=337 treatment-naïve patients. NOTCH-pathway activation and *CIITA* expression levels were inversely correlated.

Clinical impact for cases with high NOTCH-pathway activation (averaged expression levels of *HES1/2, HEY1/2*, expression above median expression level was defined as high NOTCH-pathway activation) and corresponding high or low *CIITA* expression levels (median high vs. median low *CIITA*) was assessed using gene expression data generated from PBMCs in a cohort of fludarabine resistant CLL patients.

### Statistical analyses

Data analyses were performed using GraphPad Prism 9.0 (GraphPad Software, La Jolla, USA) with unpaired or paired analyses as indicated. For experiments where more than two groups are compared, statistical analyses were performed using one-way ANOVA followed by two-tail Student t-tests. Statistical annotations were denoted with asterisks as follows: ****P < 0.0001, ***P < 0.001, **P < 0.01, *P < 0.05, and not significant (ns) P > 0.05.

