## Supplementary material for "NOTCH1 drives immune-escape mechanisms in B cell malignancies": Suppl Figure 1

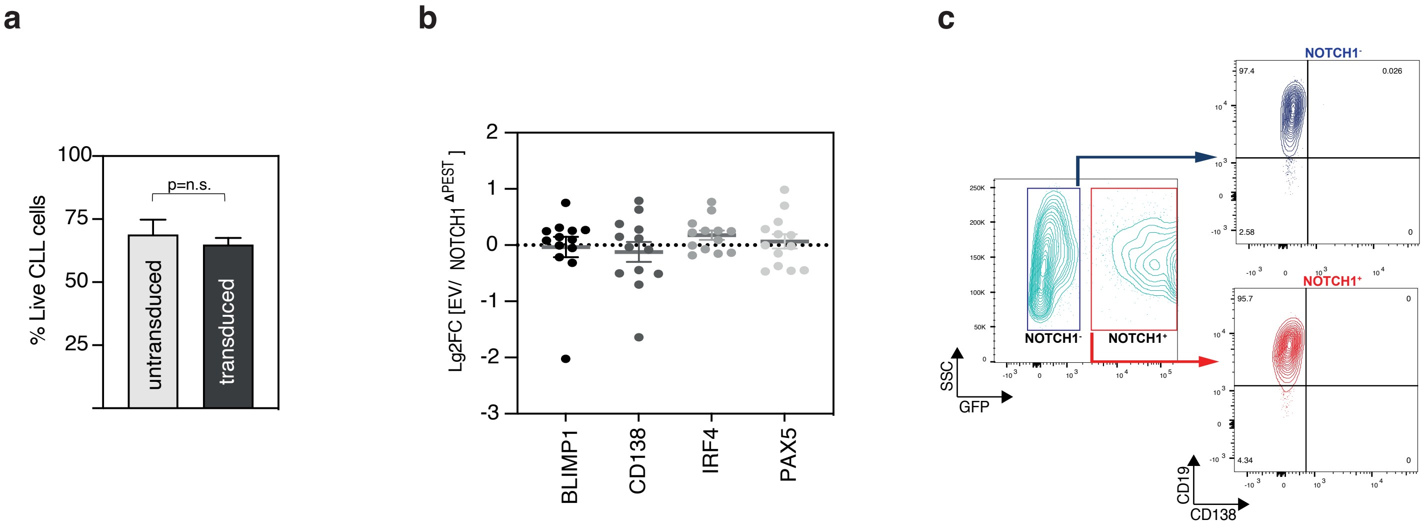


### Supplementary Figure 1:

**a**. Comparison of viable cells between untransduced and transduced CLL cells cultured under identical conditions. Apoptosis was measured by Annexin V/DAPI staining 3 days post infection [6 days post thawing].
Shown is the mean ± SEM of five independent experiments with individual primary CLL cells. ns=non significant

**b**. Log2FC values of genes associated with plasma cell differentiation, analyzed by RNAseq following NOTCH1$^{\Delta PEST}$ transduction.

**c**. CD138 expression on CLL cells transduced with NOTCH1$^{\Delta PEST}$ (red) compared to untransduced cells (blue) 5 days post infection.

**Mangolini et al.**
**Supplementary Figure 1**
