## Supplementary material for "NOTCH1 drives immune-escape mechanisms in B cell malignancies": Suppl Figure 2

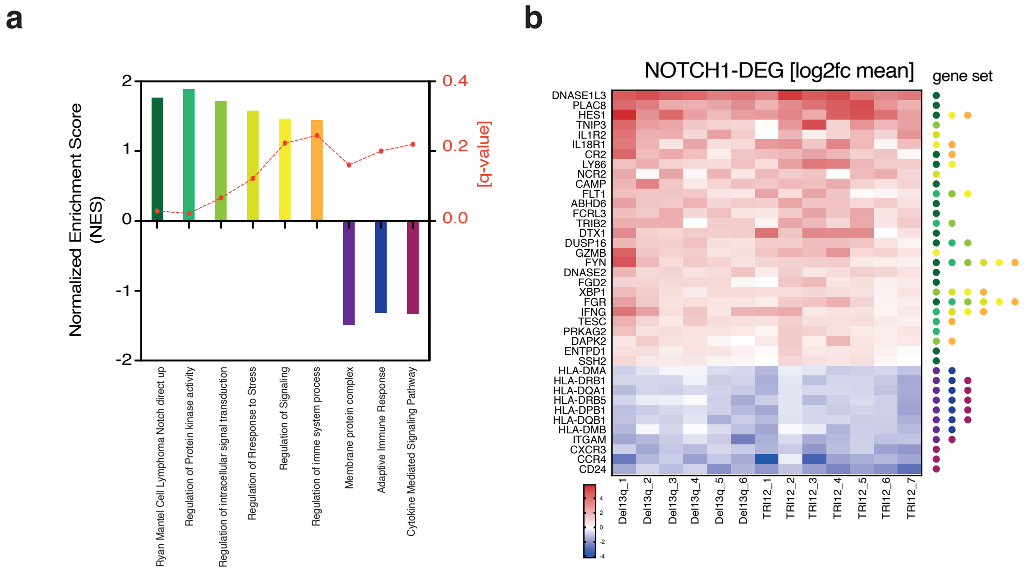


### Supplementary Figure 2:

**a.** Gene set enrichment analysis (GSEA) results of NOTCH1$^{\Delta PEST}$ genes commonly DE in Trisomy 12 and Del13q primary CLL cells.

**b.** Heatmap of the core genes of the datasets identified in A. The identified dataset to which each gene belongs to is shown using a color-coded system, indicated on the right.
