## Supplementary material for "NOTCH1 drives immune-escape mechanisms in B cell malignancies": Suppl Figure 3

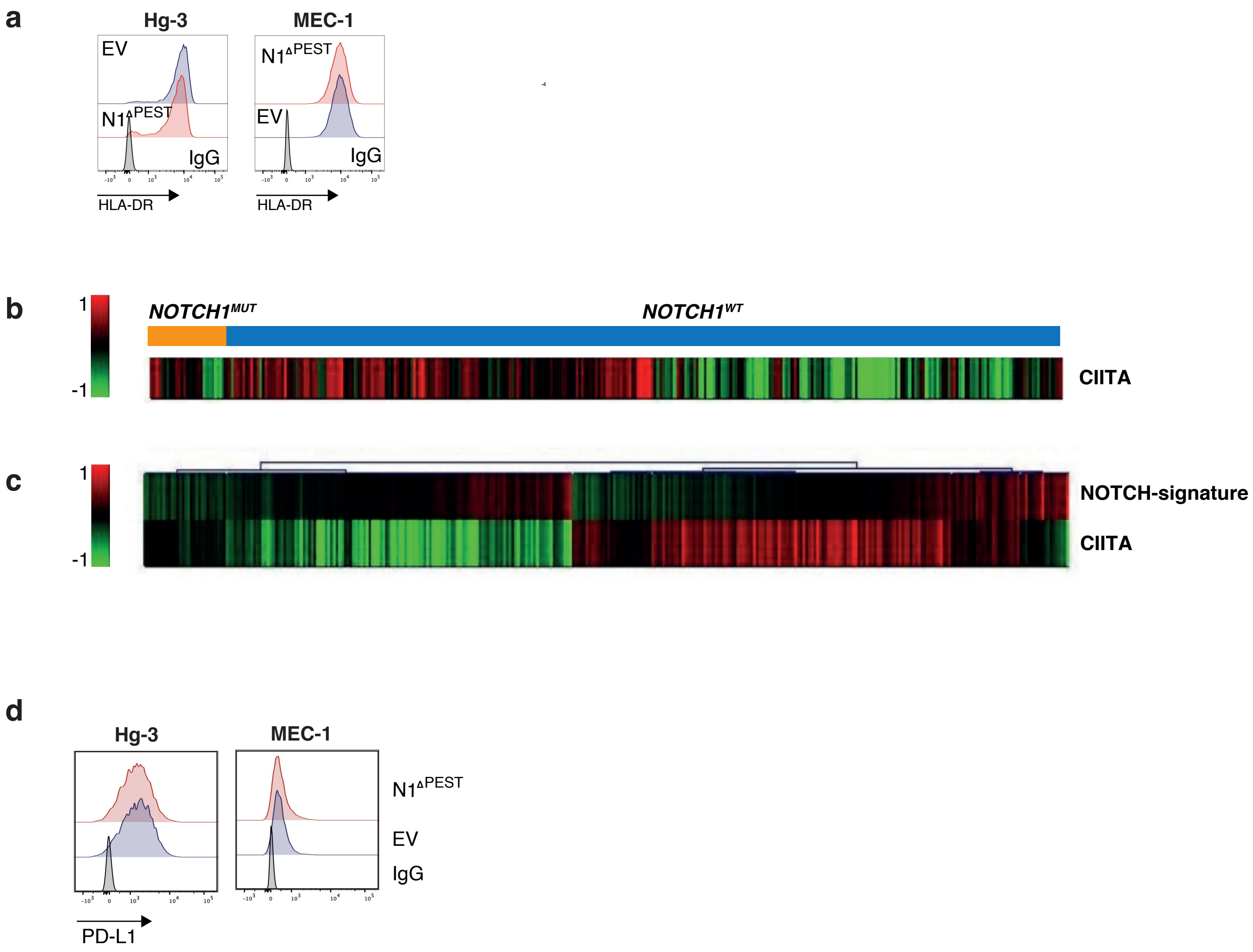


### Supplementary Figure 3:

**a.** Flow cytometry analysis of HLA-DR expression on CLL cell lines (Mec-1 and Hg-3) following expression of NOTCH1$^{\Delta PEST}$. One representative experiment out of three is shown.

**b.** Heatmap showing *CIITA* expression in treatment-naïve *NOTCH1*-mutated and wild-type CLL patients (n=337).

**c.** Heatmap of mRNA expression profiles of the averaged expression levels of *HES1/2*-,*HEY1/2*- and corresponding *CIITA* levels in treatment naïve CLL (n=337).

**d.** Flow cytometry analysis of HLA-DR expression on CLL cell lines (Mec-1 and Hg-3) following expression of NOTCH1$^{\Delta PEST}$. One representative experiment out of three is shown.
